# OCTAD: an open workplace for virtually screening therapeutics targeting precise cancer patient groups using gene expression features

**DOI:** 10.1101/821546

**Authors:** Billy Zeng, Benjamin S. Glicksberg, Patrick Newbury, Jing Xing, Ke Liu, Anita Wen, Caven Chow, Bin Chen

## Abstract

One approach to precision medicine is to discover drugs that target molecularly defined diseases. Voluminous cancer patient gene expression profiles have been accumulated in public databases, enabling the creation of a cancer-specific expression signature. By matching this signature to perturbagen-induced gene expression profiles from large drug libraries, researchers can prioritize small molecules that present high potency to reverse expression of signature genes for further experimental testing of their efficacy. This approach has proven to be an efficient and cost-effective way to identify efficacious drug candidates. However, the success of this approach requires multiscale procedures, imposing significant challenges to many labs. Therefore, we present OCTAD: an open workplace for virtually screening compounds targeting precise cancer patient groups using gene expression features. We release OCTAD as a web portal and standalone R workflow to allow experimental and computational scientists to easily navigate the tool. In this work, we describe this tool and demonstrate its potential for precision medicine.

## Introduction

Many cancers are understudied because they are rare or of little public interest, such as Ewing sarcoma, a rare pediatric cancer ^1^, and hepatocellular carcinoma (HCC), a common adult malignancy in Asia, but an orphan disease in the U.S ^2^. As the field of precision medicine progresses and we start to tailor treatments for cancer patients classified by their clinical and molecular features such as *MYC* amplification and *PIK3CA* mutation, an increased amount of cancer subtypes is emerging. The effect of each understudied cancer or cancer subtype in healthcare may be limited, but the cumulative effects of all these diseases could be profound. Ewing sarcoma is one of over 6000 rare diseases in the U.S., affecting about 2.9 per million^1^, while all rare diseases affect an estimated 25 million people in the U.S^3^. HCC affects less than 200,000 people in the U.S but is the cause of half a million deaths annually worldwide ^2^. One common research challenge for these diseases is that the resources allocated to them are relatively limited. Compared to common conditions, the large-scale screening of compounds is often challenging, if not impossible, to perform in small labs due to limited resources. The decreasing cost of sequencing enables the generation of gene expression, such as RNA-Seq, profiles of understudied cancer patient samples. Integrating these profiles with the increasing amount of other available open data (BOX 1) provides tremendous opportunity to computationally identify new potential therapeutic candidates.

Like many other investigators ^4–9^, we utilized a systems-based approach that employs gene expression profiles of disease samples and drug-induced gene expression profiles from cancer cell lines to predict new therapeutic candidates for HCC ^7^, Ewing sarcoma ^8^, and basal cell carcinoma ^9^. A disease signature is defined as a list of differentially expressed genes between disease samples and control samples (i.e., normal tissues). The essential idea is to identify drugs that reverse the gene expression signature of a disease by tamping down over-expressed genes and stimulating weakly expressed ones. In the Ewing sarcoma study, this systems-approach achieved a hit rate of >50% in predicting effective candidates ^8^. In the HCC study, we identified deworming pills as therapeutic candidates for HCC and demonstrated that the expression of disease genes was reversed in a clinically relevant mouse model after drug treatment ^7^. The recent pan-cancer analysis demonstrated that the reversal of cancer gene expression correlates to drug efficacy ^10^. Compared to the commonly used target-based drug discovery approach that focuses on interfering with individual targets, this systems-approach aims to target a list of critical features of the disease (Figure S1). The previous studies suggested that this efficient and cost-effective approach could be explored to virtually screen novel compounds or repurposing drugs using existing drug libraries such as LINCS L1000 ^10–12^.

We have shown that the success of this approach is made possible by multiscale procedures, such as quality control of tumor samples, selection of appropriate reference normal tissues, evaluation of disease signatures, and integration of drug expression profiles from multiple cell lines. For example, the scarcity of adjacent normal tissues for many cancers (e.g., pediatric brain cancers) prevents the creation of disease gene expression signatures using traditional methodologies ^13^. In TCGA, an adult cancer genomic database, less than half of cancers have at least 10 adjacent normal tissues. In the pediatric cancer genomic database TARGET, none of cancers have at least 10 adjacent normal tissues. Among these tumor tissues, a substantial number of tissues are impure, leading to a significant bias in the subsequent genomic analysis, including disease signature creation ^14,15^. There is a plethora of relevant datasets and analysis modules that are publicly available, yet are isolated in distinct silos making it tedious or even impossible to implement this approach in translational research for many labs. In this work, we describe in detail how these resources and data types can be used as well as challenges with the process. Further, we introduce our publicly-available framework and workflow, the Open Cancer TherApeutic Discovery (OCTAD), to streamline the various computational tasks required for the drug discovery. We make OCTAD available both as a standalone software package in R for bioinformaticians as well as a web server resource for investigators without a coding background. We further go through sample protocols detailing the power of our approach and the novel aspects that enable more refined prediction methods. We demonstrate the consistency of the results between the new version and our previously published HCC work, and the feasibility of using the new version to predict candidates for *MYC* amplification lung adenocarcinoma and *PIK3CA* mutation breast cancer.

### BOX 1: Public data sources and repositories

Results from laboratory experiments push scientific knowledge forward, but the raw data generated are also of particular importance. By releasing raw data into an open repository, scientists break their work out of a silo and facilitate further research and reproducibility efforts ^1,2^. For instance, other researchers can re-analyze or combine data from many experiments into meta-analyses not possible with each study in isolation. This is especially important for experiments involving rare diseases or uncommonly used cell or tissue types in which data are scarce. Here, we detail a few key open repositories for both disease and drug data.

The Gene Expression Omnibus (GEO; https://www.ncbi.nlm.nih.gov/geo/) from NCBI is a public functional genomics data repository, consisting of over three million samples from over 110,000 studies as of September 2019. ArrayExpress (https://www.ebi.ac.uk/arrayexpress/) is another functional genomics dataset that have over 55 TB of data from over 70,000 experiments as of September 2019. The Immunology Database and Analysis Portal (ImmPort; www.immport.org) is a collection of immunology-related studies with genomics and clinical outcome measurements. The Cancer Genome Atlas (TCGA; https://cancergenome.nih.gov) is a compilation of cancer-related genomics data and outcomes. The Genomics Data Commons Portal (GDC: https://portal.gdc.cancer.gov/) organizes and harmonizes TCGA data and consists of over 350,000 files from about 33,000 cases of 69 Primary Site cancers (Data Release 13.0). The Cancer Cell Line Encyclopedia (CCLE; https://portals.broadinstitute.org/ccle) details genetic and pharmacologic properties of human cancer cell line models. As of November 2018, CCLE has data for 1,457 cell lines comprised of over 136,000 data sets. Met500 is a resource profiling whole exome and transcriptome data from 500 adult patients with metastatic solid tumors of various lineages and biopsy site^16^. The Treehouse Childhood Cancer Initiative (https://treehousegenomics.soe.ucsc.edu/) is a resource that collects and distributes genomic and clinical data related to childhood cancers and contains over 11,000 tumor samples. The Genotype-Tissue Expression (GTEx; https://gtexportal.org/home/) contains genotype and expression data for almost 12,000 samples across 53 tissues from over 700 healthy donors (version V7).

For drug-related data, Connectivity Map (CMap; https://www.broadinstitute.org/connectivity-map-cmap and https://clue.io/cmap) from the Broad Institute is a large database of chemical perturbations on various cell lines. Specifically, CMap contains data on transcriptional expression changes due to administration of various chemical compounds on various cell lines. To scale this project up, CMap evolved into the Library of Network-Based Cellular Signatures (LINCS; http://www.lincsproject.org/) project, where a “landmark gene set” or L1000 of a 978 gene panel has been used to characterize combinations of 41,847 small molecules, and 75 cell lines.

## OCTAD pipeline

### Overview of OCTAD

The system includes four main components: OCTAD Dataset, OCTAD Core, OCTAD Desktop, and OCTAD Portal (Figure 1a). OCTAD Dataset stores all sample expressions, processed using Toil ^17^. OCTAD core includes all R functions needed for all analysis. OCTAD Desktop is an R markdown file which can run in a regular laptop. Its customized functions allow computational biologists performing more advanced analysis (Figure 1b). OCTAD portal is a front-end based on Python Flask and HTML5, supported by the back-end OCTAD core. We developed a simple four-step strategy to allow scientists without any programming skills to easily perform drug candidate predictions (Figure 1c). We opted to use Python Flask and HTML5, as the portal uses advanced features such as sample visualization, job management, and parallel computing.

**Figure 1:**
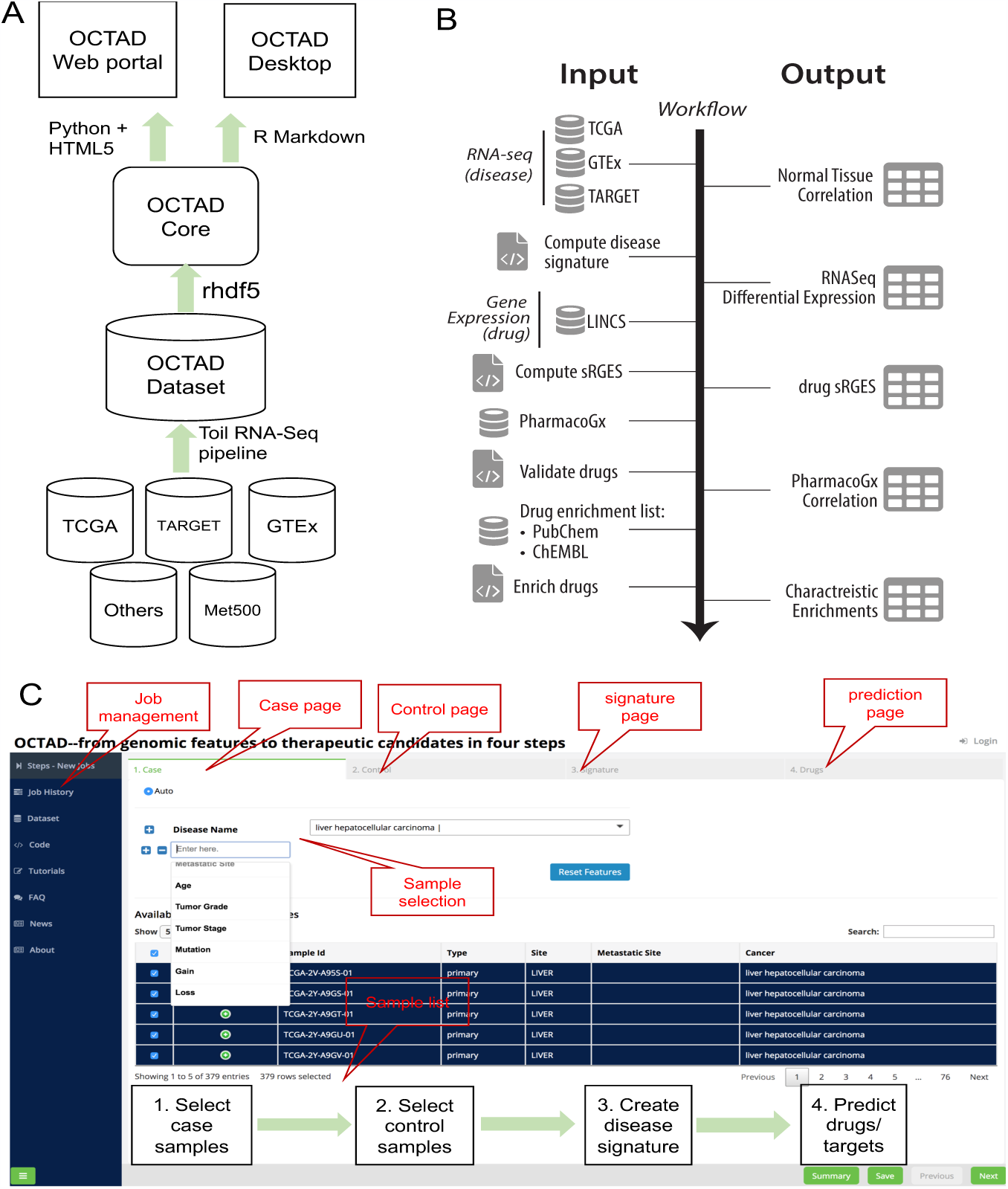
Systems description. (a) system design, (b) workflow for drug prediction, and (c) web portal screenshot.

#### OCTAD Dataset

To minimize the batch effect from multiple studies, we use the same pipeline Toil developed by UCSC to process all raw RNA-Seq profiles. We estimated transcript abundance estimated from STAR ^3^ and RSEM ^4^. Because the UCSC Treehouse initiative has already used this pipeline to process samples publicly available, we use their processed samples and extend this pipeline to process new samples. This pipeline was verified in our recent studies ^5,6^. Any new samples from the major RNA-Seq repositories including GEO, dbGAP, and EBI EGA can be easily processed by our pipeline. We have included samples from TCGA, TARGET, GTEx, and Met50, totaling 19,128 samples covering 40 cancer types (Figure 2, Table 1). When possible, we also collected their clinical (e.g., age, gender) and molecular features (e.g., mutation status) that allow the selection of a specific set of disease samples. In addition to tissue samples, we compiled 66,612 compound gene expression profiles consisting of 12,442 distinct compounds profiled in 71 cell lines (with 83% of the measurements made primarily in 15 cell lines), using data downloaded from the LINCS consortium. Each profile includes the expression measurement of 978 ‘landmark genes’. The changes in the expression of these landmark genes were computed after compounds were tested in different concentrations (62% of the measurements were made in conditions under 10 μM) for 24 h (49%) or 6 h ^10^.

**Table 1:**
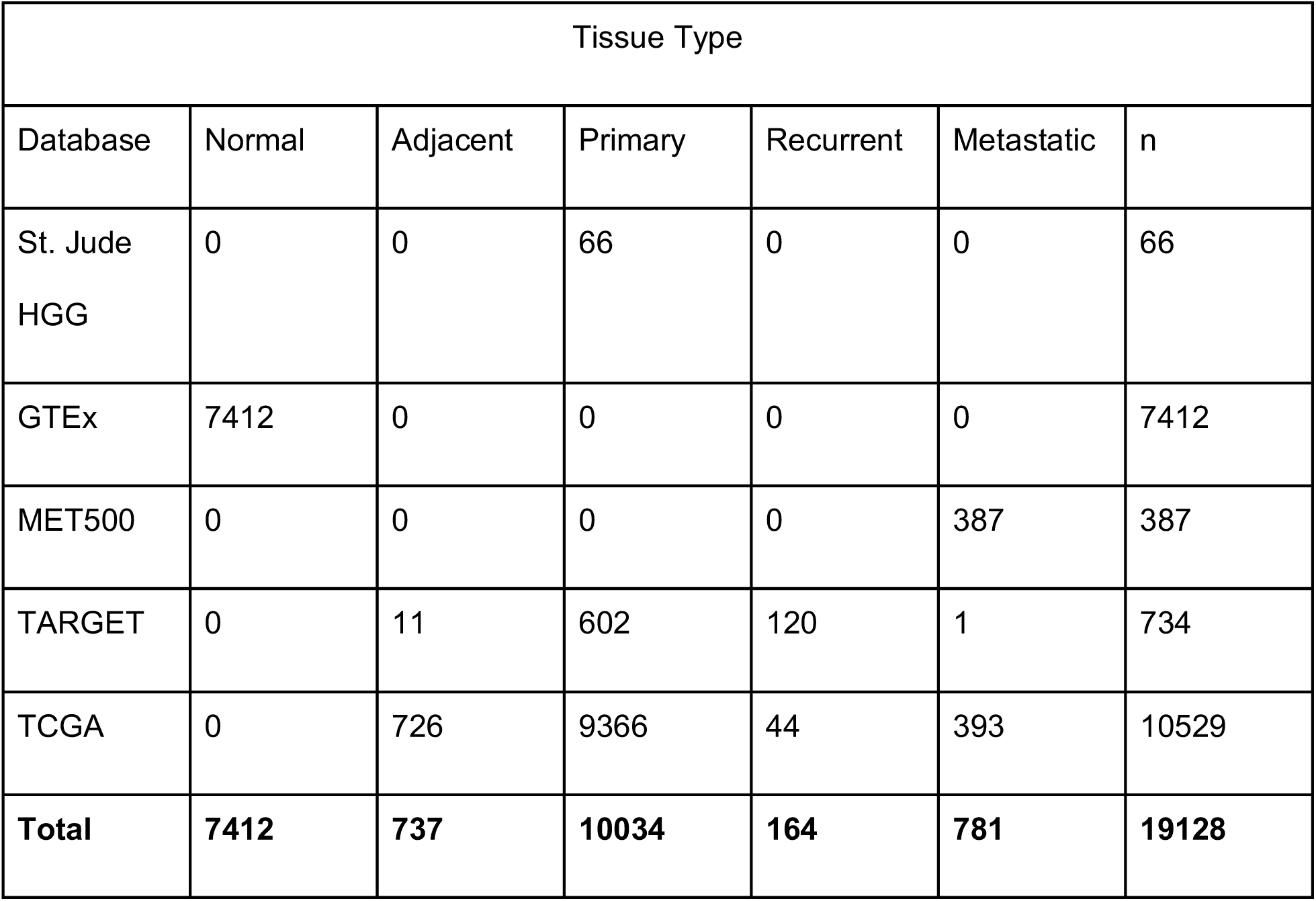
Patient sample statistics.

**Figure 2:**
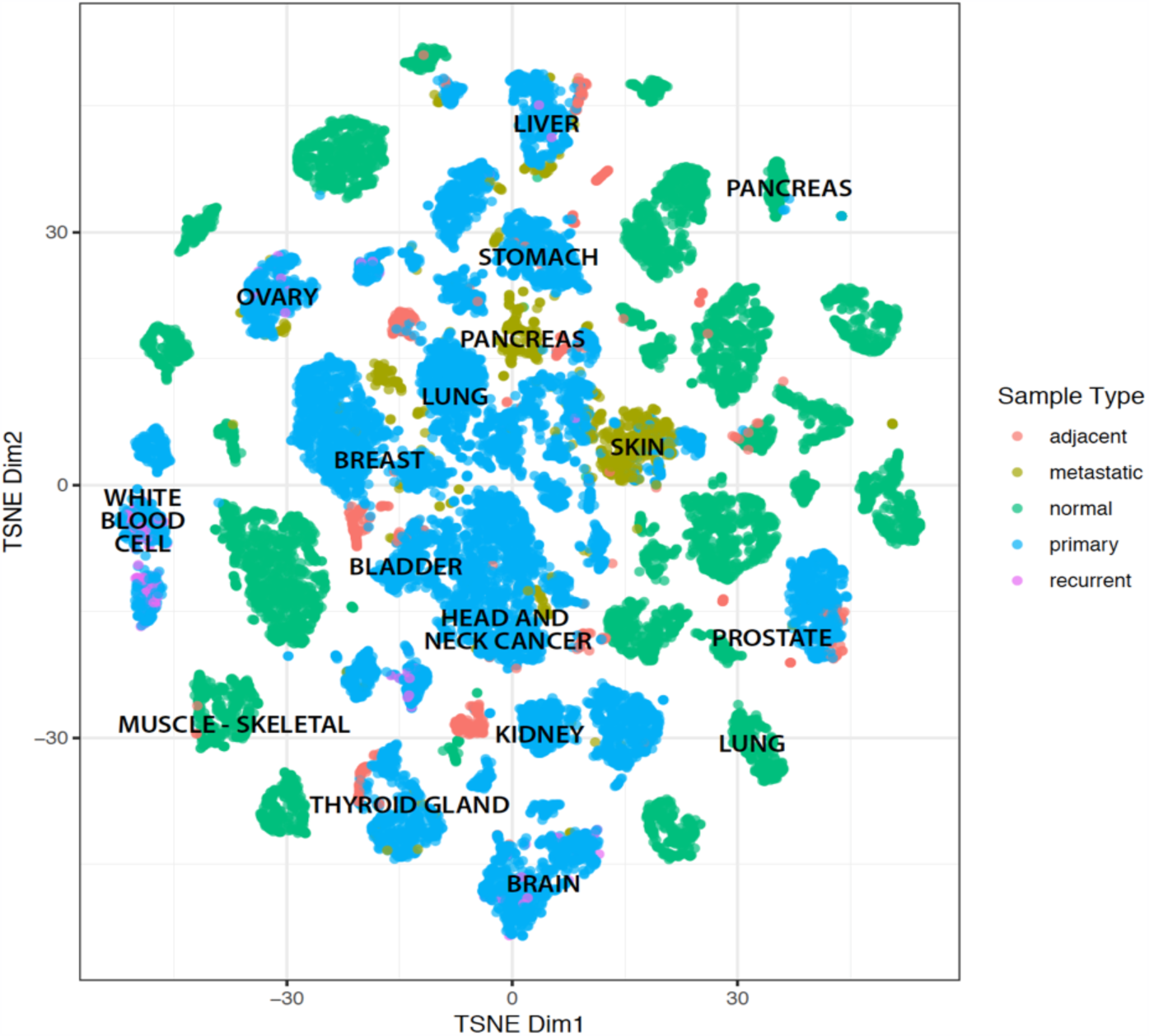
OCTAD cancer maps

#### Disease signature creation

A disease gene expression signature is defined as a list of differentially expressed genes between a specific set of case samples and matched reference normal samples. Because of the dearth of matched adjacent normal tissues, we seek to leverage normal tissue samples from GTEx, a repository of tissue samples from healthy individuals. However, the technical variation between multiple resources and the high dimensional features makes the selection of normal samples challenging. Accordingly, we utilize new features encoded by deep learning (DL) autoencoder to select highly correlated normal samples given a set of disease samples as we demonstrated in our prior work ^13^. With two groups of samples, standard differential expression analysis methods (e.g., edgeR ^18^, Limma Voom^19^) could be performed, followed by enrichment analysis using enrichR ^20^.

#### Reversal of cancer expression

In our earlier studies, we quantified the reversal of disease gene expression as RGES (Reversal Gene Expression Score), a measure modified from the connectivity score developed in other studies ^4,21^. To compute RGES score, we first rank genes based on their expression values in each drug signature. An enrichment score/s for each set of up- and down-regulated disease genes are computed separately using a Kolmogorov–Smirnov-like statistic, followed by the merge of scores from both sides (up/down). The score is based on the amount to which the genes (up or down-regulated) at either the top or bottom of a drug-gene list ranked by expression change after drug treatment. One compound may have multiple available expression profiles due to having been tested in various cell lines, drug concentrations, treatment durations, or even different replicates, resulting in multiple RGES for one drug-disease prediction. Therefore, we developed a summarization method to mitigate bias and to compute a score representative of the overall reversal potency of a compound to a particular cancer. We termed this score summarized RGES (sRGES) ^10^. We set a reference condition (i.e., concentration of 10 um, treatment duration of 24 hours) and used a model to estimate a new RGES if the drug profile under the reference condition was not available. We then weighted the RGES by the degree of correlation between the gene expression profiles of the disease and the cell line in which the compound was tested. We demonstrated that sRGES is correlated to drug efficacy (measured by IC50) and such correlation is retained even when the disease is not represented by cell lines of its own lineage in the drug expression databases. The analyses suggested the feasibility of applying this approach for large-scale screening of compounds for a given disease signature.

CAUTION

Generation of sRGES using ranked genes rather than absolute magnitude is currently the superior approach. Previous attempts to use any form of absolute expression magnitude have resulted in less robust results. While computing sRGES, restricting the cell lines to the cancer of the same lineage used by LINCS L1000 causes significant loss in the number of drugs evaluated and has not been shown to improve specificity of results.

#### Hit prediction and selection

Previous drug-repositioning efforts only considered a couple of thousand FDA-approved drugs with more potential to translate into the clinic, leaving over 10,000 compounds in LINCS unused for a broad chemical space for discovery (Figure 3a). Including those unused compounds may increase the chance of discovering novel compounds. Among these compounds, 14% are commercially available in ZINC ^7^, one of the largest collections of commercially available compounds (Figure 3b). An additional 5% of compounds are structurally similar to ZINC compounds (Similarity > 0.9), leaving more than 80% compounds that are not directly purchasable (Figure 3b). According to synthetic accessibility scores ^8^, 70% of these inaccessible compounds can be easily synthesized (Figure 3c). This protocol added a number of enrichment analyses of drug hits including enriched MeSH terms, protein targets, and chemical scaffolds. MeSH pharmacological classification and protein targets of LINCS compounds were retrieved from PubChem ^9^ and ChEMBL^10^, respectively (Figure 3d,e). Chemical scaffolds of LINCS compounds are created using RDkit (see Supplementary Material) ^11^. We expect such information will facilitate the selection of representative compounds which could be quickly obtained for testing.

**Figure 3.**
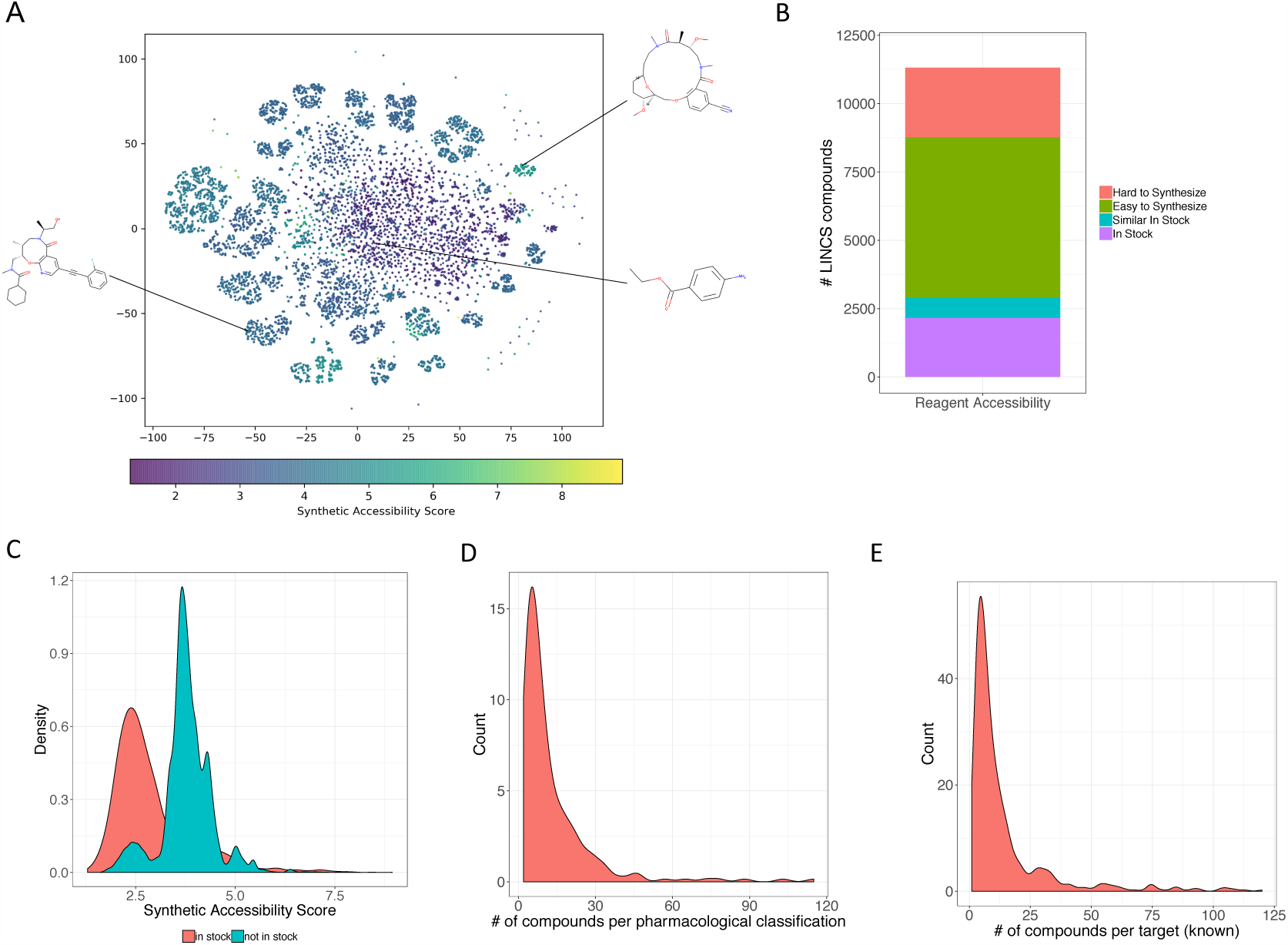
OCTAD compounds. (a) t-SNE plot of LINCS L1000 compounds colored by synthetic accessibility. This dataset covers both the traditional drug-like chemical space (purple dots) and novel chemotypes (green dots). (b) Reagent accessibility (in stock: available in ZINC, similar in stock: structurally similar to the compounds in ZINC, easy to synthesize: synthetic accessibility score < 4, and hard to synthesize: synthetic accessibility score >= 4), (c) Synthetic accessibility score distribution, (d) Mesh Pharmacological classification distribution, (e) Compound target distribution (compound targets were retrieved from ChEMBL).

#### Selection of cell lines

We rank-transformed gene RPKM values for each CCLE cell line and then ranked all the genes according to their rank variation across all CCLE cell lines. The 1,000 most-varied genes were kept as “marker genes” (we tried different gene sizes in the early preliminary analysis and did not find the large variation of results, so we decided to choose 1,000 most-varied genes in this study). Given RNA-Seq profiles of a cell line and several patient samples, we compute spearman rank correlation (across the 1,000 marker genes) between the cell line and each sample and the median value of computed spearman rank correlation values was defined as the transcriptome-similarity of the cell line with the patient samples.

#### Overview of OCTAD Desktop and Web portal

To enable users to make use of our pipeline, we release both a freely available and open source web portal and workflow in a computational pipeline. The web portal runs many of the OCTAD core functions in the backend but requires no programming expertise. It allows users to perform all parts of the pipeline including selecting case and control samples, performing differential expression analysis to generate a disease signature, and finally generating drug candidates. To make the process as efficient as possible, users can register for the web server and the various parts of the pipeline can be saved as jobs which will be saved for future visit. The web server is interactive and produces informative plots and tables that users can interact with and download. The web server also incorporates some, but not all, advanced features of the pipeline including auto-encoder recommended control sample selection. The full set of features of the web server can be found in both the Procedure section as well as the Supplementary Materials.

The desktop version can not only perform all of the above components but also provides more flexibility and features. The computational pipeline is built in the R framework and incorporates publicly available Bioconductor and R packages for processing and analyses. We provide a breakdown of the R pipeline in the Procedure section and Supplementary Materials.

### Availability

The web portal can be used by anyone without the need for extensive domain knowledge or programming expertise. Users with genetics and molecular biology knowledge will find the results highly interpretable. In order to use the desktop version, a user will need to have basic skills in R, Bioconductor packages, and knowledge of genomics data. The source codes are available in GitHub (Desktop version: https://github.com/Bin-Chen-Lab/octad_desktop; portal: https://github.com/Bin-Chen-Lab/octad_portal). The web portal is available at http://octad.org.

## Alternative Methods

The related datasets and analysis modules are publicly available yet isolated in distinct silos. Genomic data of cancer patients could be searched and visualized in platforms such as cBioPortal ^27^, Oncoscape^28^, TumorMap ^29^. Massive RNA-Seq samples are processed by platforms such as Treehouse ^17^, Rcount ^30^, and ARCHS ^31^. Disease signatures could be created by R packages such as edgeR ^18^ and DESeq ^32^. A comprehensive enrichment analysis of disease signatures could be performed in Enrichr ^20^ and DAVID ^33^. Given a disease signature, clue.io could predict drug hits using LINCS data ^12^. To be able to predict drugs using public RNA-Seq profiles, researchers have to use different platforms and utilize various tools to accomplish the task. To address this issue, we developed a portal to streamline this process. We provide an agile desktop version that allows computational scientists to customize the code, and a web portal version that allows bench scientists and clinicians to easily navigate and predict drug hits.

There are also a number of existing web resources, tools, and applications that can perform somewhat similar procedures and analyses. Rnama (https://rnama.com) is a freely available web application that allows for meta-analysis of publicly available RNA data, seamlessly extracted from GEO. Comparing case and control groups can be interactively performed which results in an interactive plot of differentially expressed genes. DEBrowser is an R shiny based package that creates an interactive dashboard to facilitate differential expression analysis and visualization of RNA count data ^34^. Users can upload their own data and the application allows for multiple QC steps and visualization. DrugSig is a web resource that allows for prioritizing potential drug repurposing opportunities, allowing for users to submit up- and down-regulated genes from pre-computed disease differential expression profiles ^35^. RE:fine drugs is an interactive dashboard that pre-calculates potential drug repurposing opportunities combining information from previously published GWAS and PheWAS results ^36^. Users can search for a drug name, disease name, or gene symbol to see suggestions based on these levels of evidence. DeSigN is an interactive web tool that allows for prediction of drug efficacy against cancer cell lines ^37^. In this application, users can enter a list of up-regulated and down-regulated genes to be compared against IC50 values to prioritize potential drugs. Drug Gene Budger is a web tool and mobile app to interactively rank drugs to modulate user-specified genes based on transcriptomic profiles ^38^. Here, users can search for a specific gene and the tool prioritizes drugs to either up- or down-regulate them using CMAP, L1000, or CREEDS data. While the aforementioned tools and applications are indeed useful, none can perform the full gambit of steps necessary for this process. Further, the flexibility of our application, the incorporation and integration of multiple datasets, along with enhancements and incorporation of novel methodologies (e.g., autoencoder reference normal sample identification) makes OCTAD truly a unique and powerful resource.

There are also many other computational resources and repositories that demonstrate the power of drug repurposing. The Drug Repurposing Hub contains an app that allows for dynamic search and exploration of annotated information pertaining to over 8,000 compounds including targets, mechanism of action, and even vendor information ^12^. RepurposeDB ^39^ and repoDB ^40^ collect and curate information about known drug repurposing experiments. These, together with studies from other researchers, demonstrate the feasibility of applying a systems approach to screen drug hits in cancers.

## Advantages, Limitations and Future Directions

Our pipeline has several advantages. First, OCTAD covers nearly 20,000 open RNA-Seq samples from multiple sources processed in the same computational pipeline so that any pipeline effect is minimized. Coupling with the robust control sample selection module makes it possible to predict drugs for cancers or cancer subtypes with no empiric controls. Second, the one-stop drug prediction web portal allows clinicians and bench scientists who may not have sufficient programming expertise to run the various computational tasks necessary to prioritize drug hits for further experimental validation. Third, the flexible desktop-based R package allows advanced users to perform customized drug discovery computation. Fourth, collected molecular and clinical samples enable precise stratification of patient samples and prediction of drug candidates for subsets of patients. Fifth, our unique deep learning-based models enable appropriate selection of normal samples. Finally, an optimal outcome is generated thanks to the rigorous quality control in each step (i.e., in silico validation of drug hits).

Despite these advantages of OCTAD, a few issues remain to address in future versions. First, the application is limited by the quality and structure of the input data, and some data sources provide more information than others, which restricts search functionality. For instance, TCGA provides a much more comprehensive coverage of clinical and molecular features than many individual studies. Similarly, these various datasets have different nomenclatures in which phenotypes and diseases are characterized. In future iterations, we will perform more intensive harmonization of these labels using common data model ontologies. For now, users will have to curate their selections based on necessity, but the application provides the framework to do so. Additionally, this pipeline is only focused on cancer. We envision this application could extend easily to other phenotypes. In future iterations of the web application, we will allow for more seamless integration of data from other sources and repositories, like GEO, SRA, EBI EGA, and Treehouse. Furthermore, OCTAD utilizes only one repurposing methodology. Last but not least, as with all drug repurposing *in silico* exploration, predictions will have to go through extensive biological and clinical validation experiments in order to verify utility and efficacy.

With the rapid advances of omics technologies, we envision that the system that we develop will greatly facilitate the use of other OMICS data (proteins, metabolites, single cells) in future therapeutic discovery. As indicated, identifying therapeutic treatments involves multiple biological systems and, as such, it is only natural that the drug discovery or repurposing process should involve multiple data types across domains. The types of data that exist to broadly assess therapeutics with phenotype are vast and cross several biological domains. These include, but are not limited to, genome, transcriptome, proteome, metabolome, epigenome, and microbiome. In the space of drug discovery and repurposing, it is important to not only look at these domains across different cell types and tissues, but also under different time points and conditions, particularly when exposed to drugs. There are also many models in which to perform such experiments, including animals (e.g., rodent, zebrafish), cell lines, organoids, xenografts, and tissues.

## Materials

For web server (http://octad.org): Developed and tested in Google Chrome.

For desktop version (https://github.com/Bin-Chen-Lab/octad_desktop):

- R v.3.5.0 or newer (https://www.r-project.org)
- RStudio (https://www.rstudio.com/)
- Download dataset from https://s3-us-west-2.amazonaws.com/chenlab-data-public/octad

Required Hardware for desktop version:

- Computer with ≥ 16GB RAM and at least 4 available CPU cores.
- Hard drive with ≥ 50GB free.
- A stable broadband internet connection.

## Procedure

**CRITICAL**

Note that an example workflow RMD and results can be found in the GitHub repository. It is highly recommended to tailor the workflow document to your needs rather than attempting to use any functions independently.

### Desktop version

We illustrate the utility of the Desktop pipeline by highlighting a use-case for HCC: we provide code and data for investigating differential expression, pathway enrichment, drug prediction and hit selection, and *in silico* validation using an external dataset. In the Code folder of the Github directory you will find hcc_aenormal.Rmd. In this workflow, we will select case tissue samples from our compiled TCGA data and compute control tissues from the GTEx data.

Note that our compiled data also contains adjacent normal TCGA HCC samples which can also serve as control tissues. However, we decided not to use them in our example to demonstrate more workflow features. Explanation of our functions can also be found in the supplement.

#### Setup

##### Download code and data

The project folder layout should include code and data. The code can be downloaded from https://github.com/Bin-Chen-Lab/octad_desktop/tree/master/Code. Data can be downloaded from https://s3-us-west-2.amazonaws.com/chenlab-data-public/octad (share in S3).

##### Install required libraries

A script to install required R packages can be found on the Chen Lab GitHub (https://github.com/Bin-Lab/octad_desktop/tree/master/Code).

##### Setup folders

The folder setup should resemble the below:

“∼/desktop_pipeline/”

“∼/desktop_pipeline/code/”

“∼/desktop_pipeline/data/”

“∼/desktop_pipeline/results/”

Variables “*outputFolder”, “dataFolder”, and “CodeFolder”* are required variables to indicate the R markdown file where the outputs should be stored, where the data is stored, and where the codes are stored, respectively. When using the provided workflow RMD, they are set automatically after a user changes the “base.folder” variable.

##### Load Data

These are the data that we compiled:

- metadata.RData: contains R elements: ‘ensemble_info’, ‘merged_gene_info’, ‘breastpam50’, ‘tsne’, and ‘phenoDF’
- CCLE_OCTAD.RData: contains all R variables necessary for cell line evaluation step
- encoderDF_AEmodel1.RData: matrix generated with AutoEncoder for control selection
- random_gsea_score.RData: precomputed random gsea scores for faster enrichment analysis with increased power
- cmpd_sets_chembl_targets.RData
- cmpd_sets_ChemCluster_targets.RData
- cmpd_sets_mesh.RData
- cmpd_sets_sea_targets.RData
- repurposing_drugs_20170327.csv: compiled list of FDA-approved drugs
- octad.h5: contains gene expression data as log2 count and log2 tpm.
- lincs_sig_info.csv
- lincs_signatures_cmpd_landmark.RData

#### Differential Expression

##### Select Case

**CRITICAL STEP**

The case_id and control_id variables must be simple character vectors containing sample IDs which match those found in metadata and expression data. They are most easily generated by subsetting the metadata matrix “phenoDF”, but advanced users may assemble them using other means.

Phenotype data contains tissue types such as normal, adjacent, primary, recurrent, or metastatic cancer. We will select for primary hepatocellular carcinoma. The code as below:

phenoDF_case <- phenoDF %>% filter(cancer == ‘Liver Hepatocellular Carcinoma’, sample.type == ‘primary’, data.source == ‘TCGA’)

case_id <- phenoDF_case$sample.id

The sample ids for these cancer will be stored into the variable *case_id*.

This code can be easily modified to select other cancers, or to select for mutation subtypes. We include mutation data in the datasets we provide, so it would be simple to filter cancer based on mutation status such as *TP53* or *MYC* mutations.

**CAUTION**

Selected cases and controls are the two most important factors in achieving the best results when using this pipeline. There are several methods included in the provided code which evaluate controls relative to cases, but there are no built-in validation steps which evaluate cases. Each group of cases needs to be evaluated individually for validity by the investigator.

##### Select Control

In this workflow, we will compute correlating normal tissues from GTEx using encoded features of our dataset and the *computeRefTissue* function as below.

normal_id = (phenoDF %>% filter(data.source == ‘GTEX’))$sample.id

The pool of normal ids which will be drawn from to compute controls will be stored into the variable *normal_id*.

Control_id <- computeRefTissue(case_id = case_id, normal_id = normal_id, expSet = dz_expr, control_size = 50)

The default output of this function is to select the 50 top samples from the normal_id with the pool that has the highest median correlation to the case samples. The sample ids for control tissues will be stored into the variable *control_id.*

Figure 4a shows the top 50 tissues highlighted in red that has the highest median correlation. The tsne distribution of case and control are highlighted in Figure 4b.

**Figure 4:**
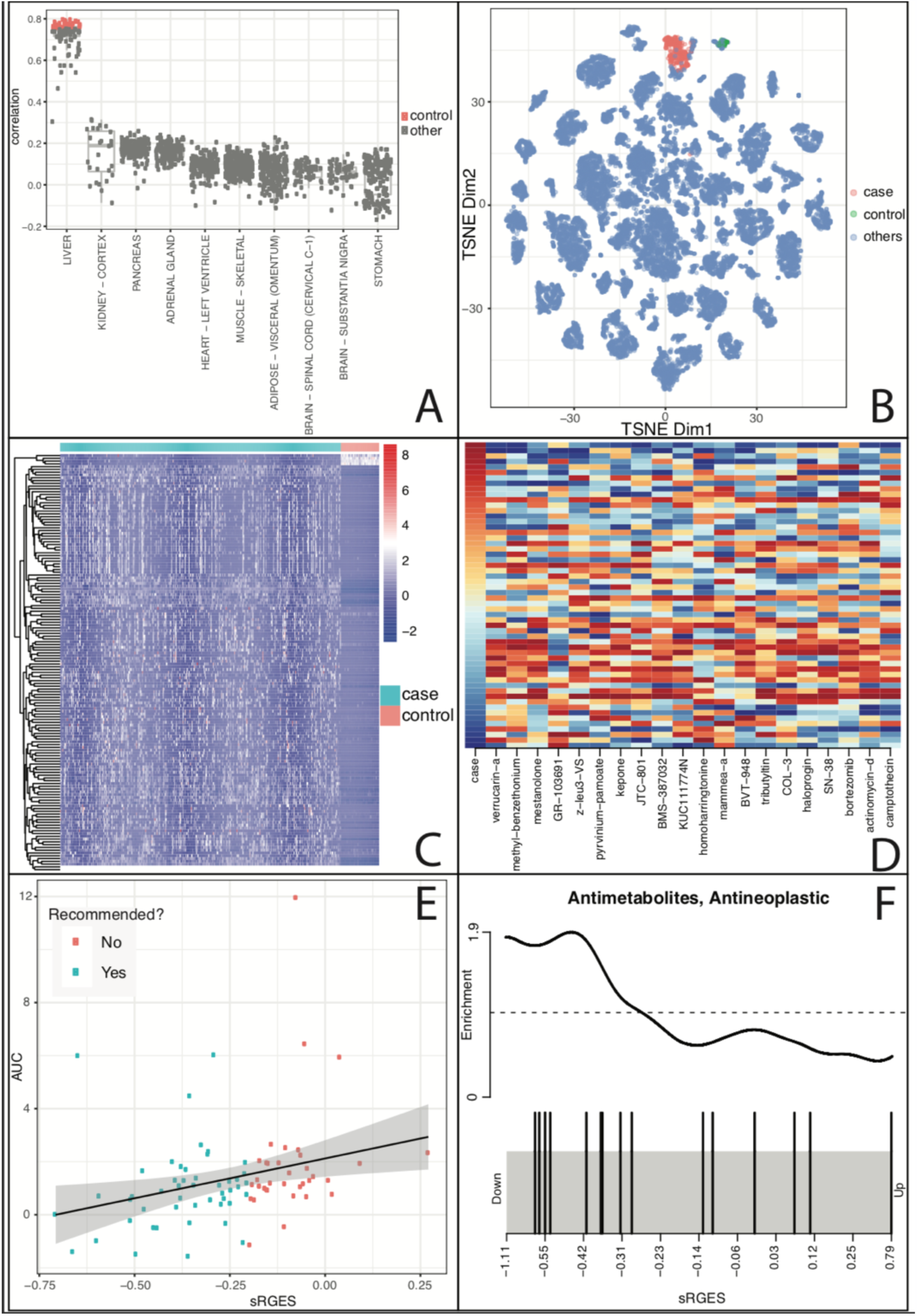
Screen compounds targeting female luminal A breast cancer. (a) correlation between luminal A breast cancer tumor samples and all samples from normal organs. Highly correlated samples (colored by red) were selected as control. (b) Distribution of selected samples in cancer map. (c) Disease signature visualization. (d) Top compounds that reverse the disease signature. (e) correlation between sRGES (predicted score) and drug efficacy data in vitro, and (f) enriched drug class. Drugs belonging to the Antimetabolite, Antineoplastic class are represented as black bars.

##### Differential Expression

Differential expression can be computed via edgeR, limma, or DESeq2. In our file we compute using edgeR. Since this function computes differentially expressed genes between *case_id* and *control_id* within the same data matrix, this can be used to find differentially expressed genes between any two groups.

res = diffExp(case_id = case_id, control_id = control_id, expSet = dz_expr, normalize_samples = T, DE_method = ‘edgeR’)

**CAUTION**

To use a pre-computed differential expression signature, make sure it follows the precise format of the example dz_signature.csv.

##### Batch Normalization

In the RMD code chunk we set *normalize_samples = T* which utilizes *RUVSeq* to mitigate batch effects ^41^. The options *k, ntop_genes*, are parameters used for *RUVSeq* and are only needed if normalize_samples is set to true. These options are used to compute an empirical set of control genes via edgeR. We suggest batch normalization when using heterogeneous samples e.g. TCGA cancers and GTEx normals as these will produce more conservative results. The disease signature is visualized in a heatmap (Figure 4c).

**CAUTION**

Default hyperparameters were found to produce the best results in a handful of validation cases. These should be evaluated for appropriateness with each new case.

##### sRGES

The *runsRGES* function is used to compute the drugs that potentially reverses the disease signature. The code below shows to filter for *significant genes* by filtering for low p adjusted values and high log fold change values.

dz_signature <- res %>% filter(padj < 0.001, abs(log2FoldChange) > 2)

runsRGES(dz_signature = dz_signature)

Note that in the runsRGES code requires use of our compiled LINCS1000 perturbation database. Since LINCS1000 only measures 978 genes, the disease signature are genes in the differential expression which overlaps with in the 978 genes of LINCS. The runsRGES will compute any dz_signature output that was generated by our diffExp function. Figure 4d is a sample output of drugs that are predicted either in clinical trials or launched.

**CAUTION**

Final results include no significance scores. Using sRGES lower than −0.2 is a method found to produce the best results in a handful of validation cases. There is currently no known superior way to compare hits to aid in lead optimization. Similarly, comparing the magnitude of sRGES results across runs is inappropriate.

##### Validate Results with CTRP (optional)

As the pharmacogenomic database CTRPv2 consists of efficacy data of 481 drugs in 860 cancer cell lines ^43^. We leverage it for further *in silico* validation of our predictions, even without running any biological experiments. We used the HepG2 cell lines to validate our sRGES list. In our previous work, we’ve shown that RGES scores correlate with drug efficacy such as AUC or IC50 ^12^. We model a simple regression to quickly validate our analysis (Figure 4e). Note that we generally use this analysis to optimize the pipeline for our disease of interest by changing various hyperparameters.

##### Drug enrichment (optional)

After calculation of sRGES on the L1000 compound dataset, drug enrichment analysis is used to summarize the result. The goal of this is to summarize and discover compounds that belong to a certain class, e.g. anti-inflammatories, EGFR inhibitors, dipines. We combined L1000 drugs into lists: MESH, CHEMBL, and CHEM_CLUSTER for MeSH term enrichment, target enrichment, and chemical structure enrichment, respectively. The enrichment score was calculated using ssGSEA ^44^. Figure 4f shows the enrichment of Antimetabolites, Antineoplastics in the prediction and Figure S2 lists these compounds.

This analysis provides much information for following candidate selection and experiment design. First, selecting candidates from the same enriched class (i.e., MeSH term, target) are more likely to be true positive candidates than random selection from the top list (Figure S2). Second, when the ideal candidate is not available, it’s reasonable to choose an alternative from the same class. Sometimes, it is necessary to choose a new drug for testing (e.g., a second generation of one inhibitor for the same target). Lastly, since many compounds have multiple MOAs, this analysis would help interpret the MOA of promising compounds.

### Web Server version

#### Portal Landing Page/Log-in

At the landing page, users can either create a new account, login to their existing account, or go straight to creating a job (Figure 5). While having an account is not required to initiate the job, we recommend setting one up as it will save all active and past jobs. User account settings can be accessed on the top right of the header. Once in the main portal, a user’s Job History can be found in the tab on the side menu. Here, all jobs are listed and include information pertaining to disease of interest, status (e.g., Completed, In Progress, etc), creation time, as well as the ability to view and download all output. Users can also delete previous jobs. At the bottom of the screen, the details of the current job can be found by clicking the Summary button, the current job can be saved clicking the Save button, and navigating forward or backwards in the process can be done by clicking the Previous and Next buttons respectively.

**Figure 5:**
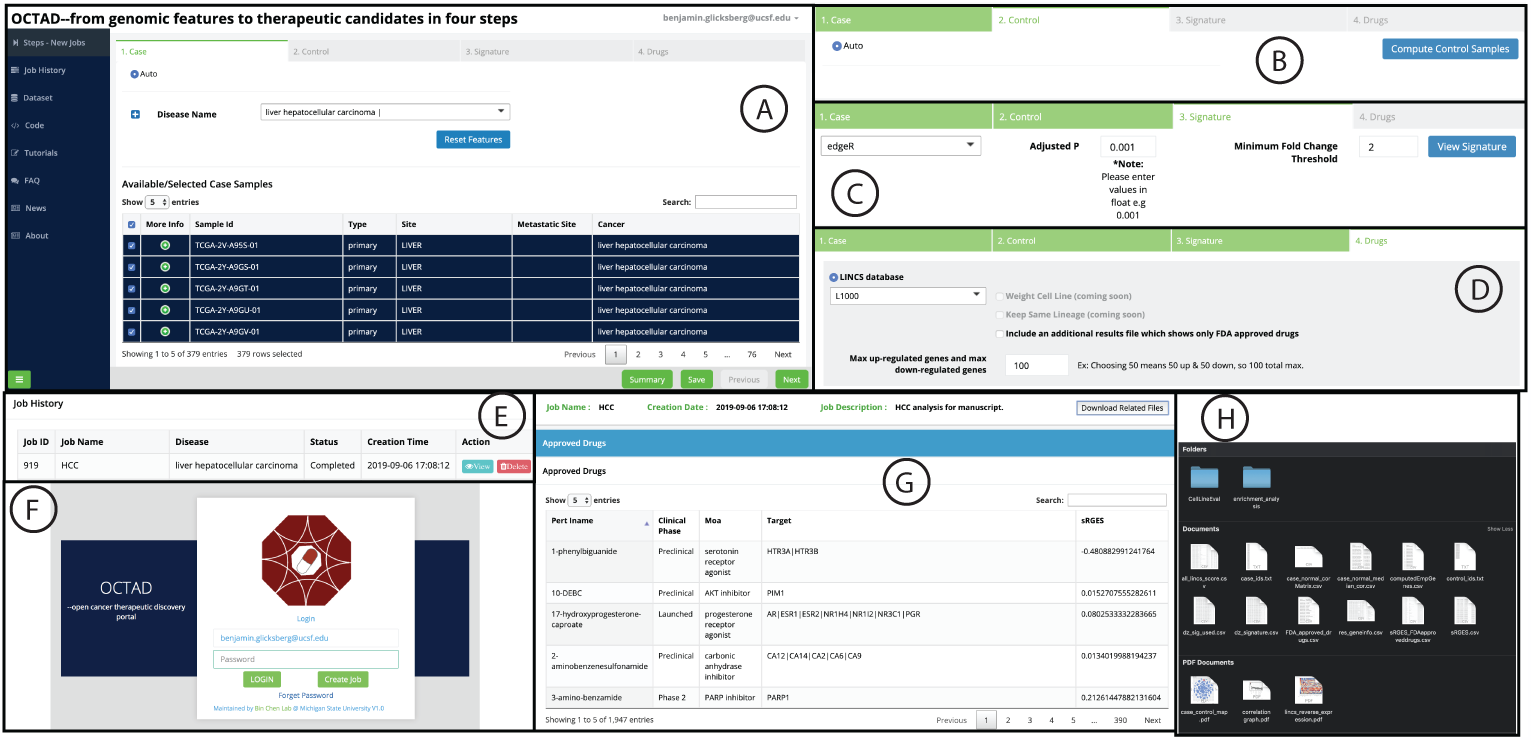
screenshots of the web portal. (a) disease sample selection, (b) control sample selection, (c) drug prediction job submission, (e) job management, (f) log in page, (g) predicted drug list, and (h) result files.

#### Creating a New Job: Hepatocellular Carcinoma

Creating a job is separated into four sections: Case, Control, Disease Signature, and Drugs. In this procedure, we will go through all steps along with expected output. We will highlight hepatocellular carcinoma as our featured example.

#### Case Selection

Upon entering the Case section, all samples available from all included database resources (see Materials; *n* =19,128) are displayed in a table at the bottom of the page, including: cancer type (Cancer), site of derivation (Site), and whether it is metastatic (Type and Metastatic Site columns). The user is able to interact with this table such as searching in the top right and viewing more information about the samples by clicking the green “+” button under More Info, which shows information like patient demographics and status for certain mutations. Users can manually select samples to include as well by selecting the check box at the left of the sample row.

To begin a job, we can search for cancers of interest in the search box at the top of the page besides Disease Name, and multiple diseases can be selected. Users are then able to further refine their search by adding filters (e.g., Gender, Tissue Type, certain mutation status), which automatically update the selected samples below. Lastly, as before, samples can be manually added and removed by checking or unchecking rows in the table respectively. We can begin our search by typing “hepatocellular carcinoma” into the Disease Name search box and choose the corresponding option. Without any other filters added, we are left with 421 samples. We can proceed to select Control samples in the next section by clicking the Next button on the bottom of the screen.

#### Control Selection

After we click compute control samples, the portal will ask for if adjacent samples should be included if available and the number of control samples should choose. The portal will recommend control samples automatically. By using the DL method as recommended, we are left with 50 samples. These Control samples are highlighted along with the Case samples from before on the box plot against all other normal samples. Depending on the user’s goals, this can be a good way to visually verify whether the current selections are adequate. We can continue to the Disease Signature section by clicking the Next button on the bottom.

#### Disease Signature Generation

With the Case and Control samples selected, we can now create the disease signature of interest. This section allows for multiple methods to accomplish this, specifically using either edgeR or limma. For this job, we will select edgeR and press Compute to begin. As indicated by the warning pop-up, this task can take a few minutes, so one benefit of creating an account is that the user can log out while this process is occurring (and even get notified by e-mail when the task is complete.). Once the signature generation is finalized, this section produces a heatmap of differentially expressed genes as well as a table below with these data. The criteria metrics can be set and changed here, specifically the p-value and fold change cut-offs. Changing these will affect the resulting plots and tables. This heatmap is interactive and can support sorting by each axis. The heatmap and table can be exported for reference.

In addition, we also integrate other tools and datasets to help make sense of the disease signature. For instance, we provide Gene Ontology pathway enrichments for both up- and down-regulated genes in the signature as both table and bar plots. With the disease signature generated, we can proceed to the final stage of the job to calculate the RGES for all available drugs by clicking the Next button on the bottom panel.

#### Candidate Drug Selection

In this section, we can compare the disease signature to all drug signatures obtained from the LINCS resource.

#### Job submission and tracking

After submitting the job, the pipeline typically takes 10-20 minutes to run. Once it is finished, a notification email will be sent to the user’s email box. Users can also monitor job status through Job History. The results files are the same as those generated from the pipeline.

## Troubleshooting

Please include information on how to troubleshoot the most likely problems users will encounter with the protocol.

Desktop Version

https://docs.google.com/spreadsheets/d/1Dy8HkC4zu50QUiAjxd-8P1daVRamQZJD3bQj2F3Ykts/edit?usp=sharing

Web Portal Version

[Table with Error messages and their meanings]

## Results

In the procedure for the Desktop version, we demonstrated the ability of the OCTAD pipeline to select a case of primary cancers from TCGA, compute correlated reference tissues from GTEx to be used as control samples, and compute a differential gene expression signature to recommend candidate drugs. This current pipeline is a complete framework with increased functionality compared to our original iteration. For instance, instead of using DESeq to compute differential expression, we adopted a less time-consuming method edgeR. Further we integrated our methodology to select reference tissue from GTEx data, making possible the prediction of cancers without adjacent normal tissues. Lastly, we compiled more samples along with their clinical features, enabling prediction of candidates for a subset of patient samples. Here, using HCC as an example, we first demonstrate the consistence of results between the original method and the optimized one in every major step (i.e., disease signature creation, drug prediction, drug enrichment analysis). To illustrate the power of using OCTAD to screen compounds for identifying putative personalized therapeutics, we predict compounds specifically targeting *MYC* amplified lung cancer and *PIK3CA* mutant breast cancer.

### Comparison between Original Method and Current Pipeline

We first compared the rank of differential expression log2 fold change from our original work which utilizes DESeq2 (noted as Pub) to the rank of differential expression in our current pipeline which utilizes edgeR (noted as Adj); both utilize TCGA adjacent normal HCC as a reference control. Next, we compared the original results with our current pipeline using edgeR and computed GTEx normal tissues derived using top correlated autoencoder method (noted as AE), generated from the desktop procedure section. Finally, we compared the results with those derived from our online web portal (noted as Online) which also utilizes edgeR and normal liver tissue from GTEx database which was generated from the online procedure section.

The correlation analysis of the differential expressions from different procedures shows that the workflow utilizing edgeR and adjacent tissues as control had the highest correlation to the original work (Figure 6a). In addition, both the online and desktop disease signatures are equal to each other as the parameters for them are the same (Figure 6a). Both online and desktop results utilizing GTEx as controls also retained high correlation to the original gene expression signatures, indicating that it’s feasible to use GTEx controls (Figure 6a).

**Figure 6:**
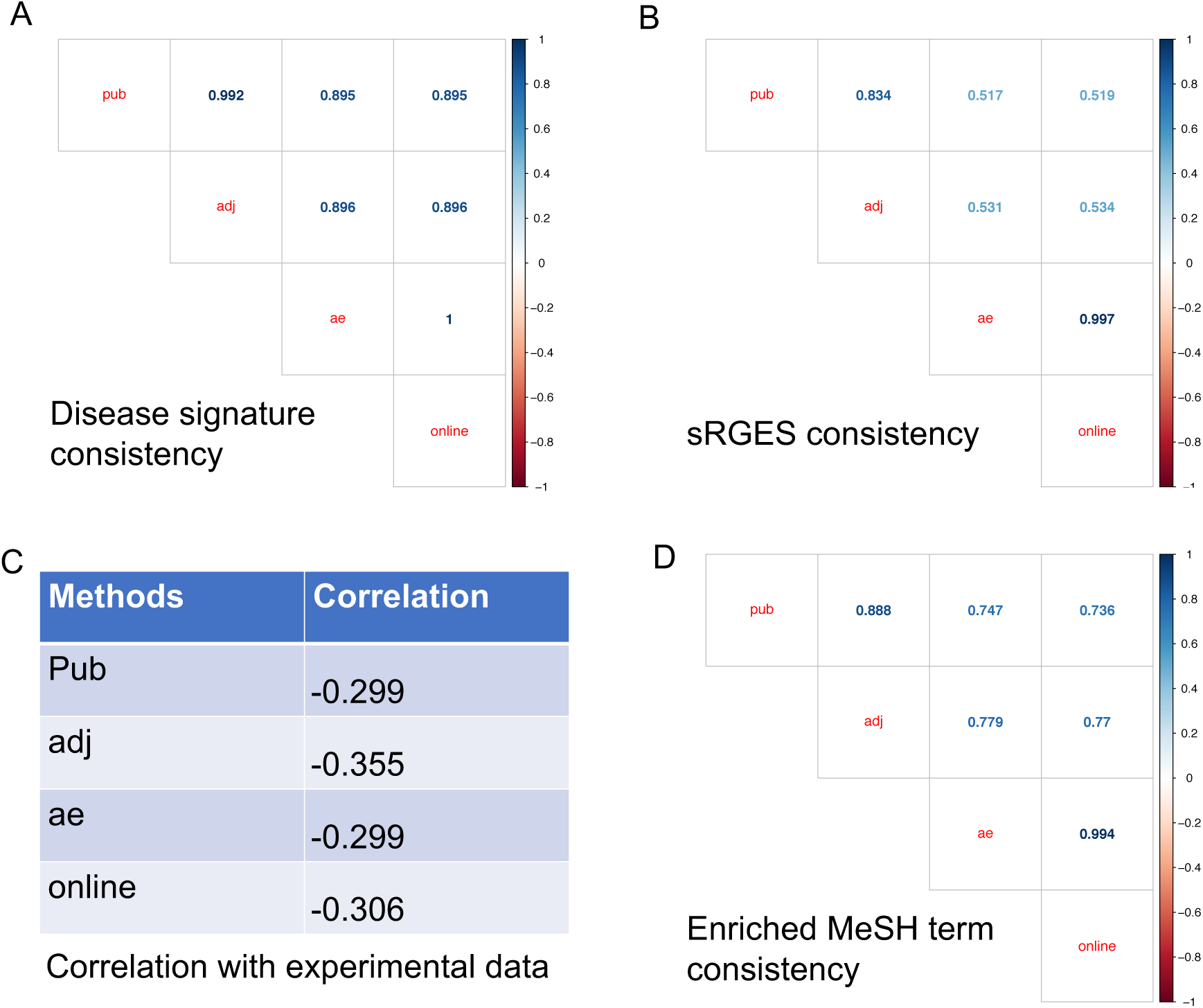
Evaluation of the results from major steps in HCC prediction. (a) consistency of disease signatures, (b) consistency of sRGES, (c) correlation with experimental data (IC50), and (d) consistency of enriched MeSH. Pub: the published work ^10^; adj: using edgeR and adjacent tissue as control; ae: using edgeR and normal tissues selected from the AutoEncoder approach; online: online web portal.

As subsequent drug prediction sRGES is computed using significant differential gene expression and we observed the little discrepancy of differential gene expression computed from multiple procedures, we then assessed if sRGES are also similar to the original. Unsurprisingly the workflow utilizing edgeR and adjacent tissues as control had the highest correlation to the original work. Both online and desktop results had lower but still significant correlation to the original work (Figure 6b).

In our original work, we showed that sRGES correlated with drug efficacy data from CTRP database in which drugs were tested on cancer cell lines. We further compared the correlation of sRGES to the AUC of the corresponding drugs found in CTRP of liver cancer lineage. AUC is computed as the median AUC across all the liver cancer cell lines. The significant correlation between drug efficacy data and sRGES computed from different procedures suggested the utility of using the workflow to *in silico* predict drug efficacy, even in the absence of adjacent tissue samples (Figure 6c).

One of the pitfalls of using individual scores such as sRGES is the possibility for false positive and false negative predictions. An additional function of our workflow includes enrichment of drug targets such as via MESH terms. This allows for summary of drugs into classes to allow for investigation of groups of drugs rather than individual compounds. A higher score indicates a MESH drug group to be efficacious against the cancer, whereas more lower and negative score indicates non-effective MESH drug group. We further compare the rank correlation of the MESH scores generated from the different procedures (Figure 6d).

By summarizing the sRGES in classes of drugs we are able to examine clusters of drugs based on their mechanism of action e.g. MESH. Finding novel classes of drugs, e.g. anti-helminths, or unconventional chemical structures allows for generation of hypotheses that can be prioritized for experimentation. Furthermore, the pipeline also allows us to filter out for drug candidates which may have low value, for example, poor scores were given for anti-hypertensives compared to more conventional classes such as intercalating agents and HDAC inhibitors.

### Screening compounds targeting *MYC*-amplified lung adenocarcinoma

Targeting MYC, an oncogene amplified in many cancers including lung adenocarcinoma, is under active research, however, MYC is considered as an undruggable target ^45^. One therapeutic strategy is to reverse the gene expression signature of these *MYC*-amplified tumors. Here, we ran the two sets of TCGA Lung adenocarcinoma through our pipeline. In one set we selected for cancer tissues with *MYC* amplification, defined as copy numbers of 1 or more. In the second set we selected for cancer tissues without the *MYC* amplification, defined as 0 copy numbers. Then we selected drug efficacy data from the CTRP non-small cell lung cancer cell lines with the *MYC* amplification as a validation set. The correlation for the MYC run was −0.415, while correlation for non-MYC run was −0.261 (Figure S3). This suggests that OCTAD may be used to search for candidates specifically targeting *MYC*-amplified cancers.

### Screening compounds targeting *PIK3CA* mutation in breast cancer

Likewise, we applied OCTAD to screen compounds targeting tumors harboring *PIK3CA* mutation. *PIK3CA* is highly mutated in cancers, yet undruggable ^46^. In this case we ran the two sets of TCGA Breast cancers through our pipeline. In one set we selected for cancer tissues with *PIK3CA* mutation. In the second set we selected for cancer tissues without the mutation. Then we selected for CTRP breast cell lines with the *PIK3CA* mutation to use as a validation set. Similarly to the previous result, we found that mutated breast *PIK3CA* cancer correlates with corresponding CTRP cell lines better than non-mutated *PIK3CA* breast cancer. The PIK3CA mutant run has correlation of −0.370, with the non-mutant run at −0.273 (Figure S4).

## Supporting information

main

## Summary

The combination of clinical and molecular features could lead to numerous disease subtypes, each of which may have limited resources to study. Through leveraging open datasets and advanced machine learning methods, OCTAD aims to provide an effective means to screen compounds for one specific cancer subtype for further experimental testing. In this work, we optimized our pipeline and developed OCTAD, which we demonstrate can reproduce results from a previous study that are also consistent when selecting normal tissue data from a different database. Furthermore, the two new cases illustrate the potential of using OCTAD to screen compounds for a precisely defined patient groups, although subsequent experimental testing is desired to verify drug candidates.

## Contributions

BZ led the project, developed the desktop version, and wrote the manuscript. BG and PG co-led the project, coordinated web portal development, and wrote the manuscript. JX performed the LINCS compound analysis and developed compound enrichment analysis. KL implemented the Toil pipeline and proceed RNA-Seq samples. AW helped develop the code, prepare tutorials and create case studies. CC helped trouble shooting. BC supervised the project.

## Acknowledgements

The research is supported by R01GM134307, R21 TR001743 and K01 ES028047 and the MSU Global Impact Initiative. The Amazon AWS research credits were received to support portal development and hosting. The portal was developed with the help from Optra Health Inc. The content is solely the responsibility of the authors and does not necessarily represent the official views of sponsors.

