## Supplementary material for "OCTAD: an open workplace for virtually screening therapeutics targeting precise cancer patient groups using gene expression features": main

### Supplementary text

**Clustering.** All the compounds in LINCS L1000 database (11313 compounds with known chemical structures) were clustered into thousands of compound groups based on their chemical structural similarities with each other. The similarity score of two compounds was calculated as the tanimoto coefficient of their 2048-bit FCFP8 fingerprint pair by RDKit. The Butina Algorithm in RDKit was employed to perform the chemical structural clustering. As the L1000 molecule dataset is highly diverse and covers a large chemical space of both traditional drug-like molecules and novel complicated scaffolds, a simple threshold of intra-cluster distance was not competent to this task. Thus, different intra-cluster distances from 0.4 to 0.8 were used to gather the molecules at different levels. All compounds were first attempted to be clustered with a cutoff of 0.4, then those classes with only one or two members were re-clustered with a cutoff of 0.5. These steps were repeated until a cutoff of 0.8 was achieved. Finally, compounds with similar chemical structures (or substructures) were grouped together.

**Enrichment analysis.** After structural clustering, compounds belonging to each group were calculated enrichment using ssGSEA. The p value was derived from the frequency of lower ssGSEA scores of the 1000 randomly altered sRGES rankings. A default p value of 0.05 was set for further analysis.

**Purchasability fetching.** All the compounds in LINCS L1000 database were searched for purchasability in ZINC “in stock” subset of compounds, including the exact structure searching and similar structure (tanimoto coefficient > 0.9) searching.

**Synthetic accessibility scoring.** All the compounds in LINCS L1000 database were estimated their synthetic accessibility by the SA_score in RDKit. This score ranges from 1 (easy to make) to 10 (very difficult to make).

**Distributions of chemical clustering, purchasability and synthetic accessibility of L1000 compounds.**

In this research, 2695 compound groups were obtained after clustering. Among them, 2078 groups contained no less than 3 members. As shown in the t-SNE plot (figure x1), L1000 dataset covers a broad chemical space. The novel complicated chemotypes (green dots) were far from the traditional drug-like molecules (purple dots). There are 2161 purchasable molecules, and 742 molecules with their analogues purchasable. About half of the L1000 compounds were predicted with low or moderate synthetic difficulty.

### Supplementary Figures

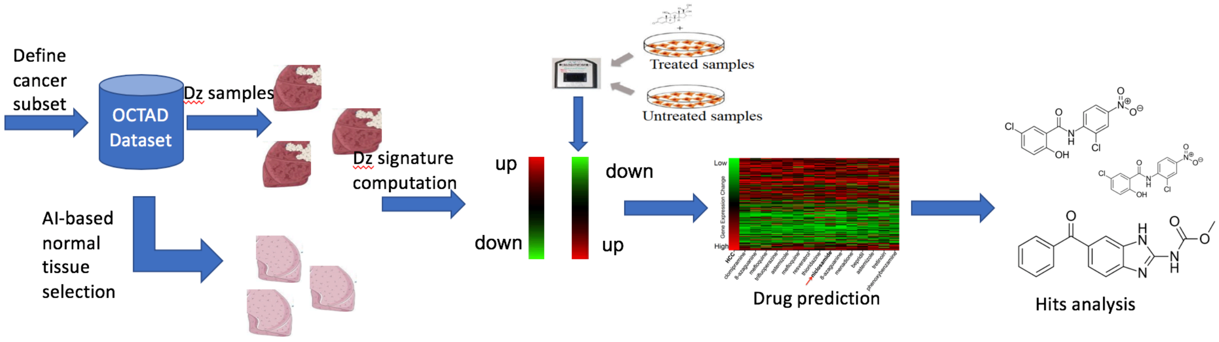


Figure S1: Illustration of they systems-based therapeutic discovery approach. AI: Artificial Intelligence; Dz: Disease

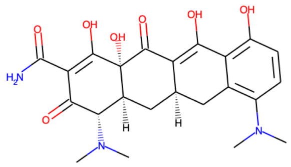

Minocycline

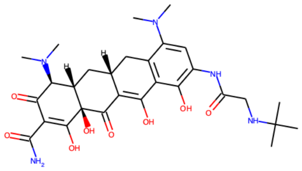

Tigecycline

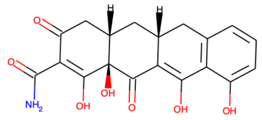

COL-3

| Compound | sRGES | Purchasable | Potency |
|---|---|---|---|
| Minocycline | -0.494 | Yes | Inhibition of metastasis combined with celecoxib |
| Tigecycline | -0.379 | Yes | IC50 10-25 µM against MCF-7 and T47D |
| COL-3 | -0.378 | Yes | Enhances chemotherapy for MDA-MB-231 and pII |

Figure S2: Example of enriched drugs

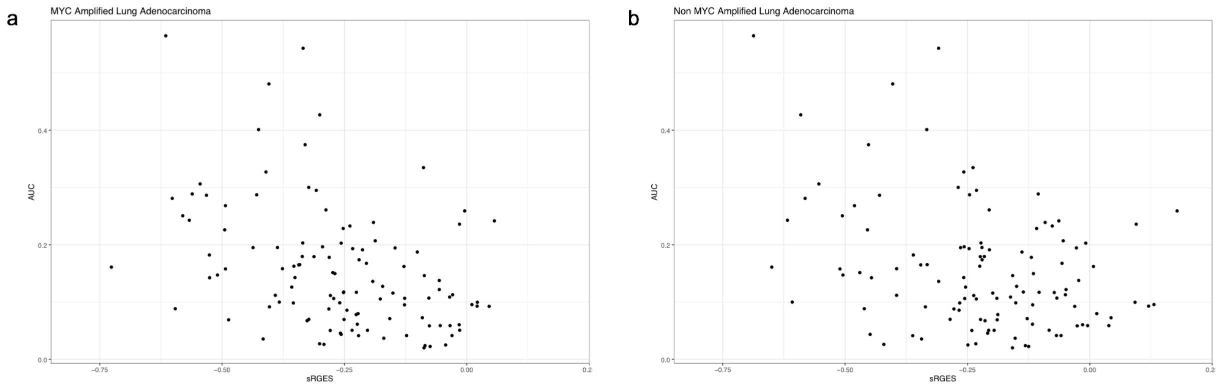


Figure S3: Correlation between sRGES and AUC in (a) MYC amplified lung adenocarcinoma, and (b) non-MYC amplified lung adenocarcinoma.

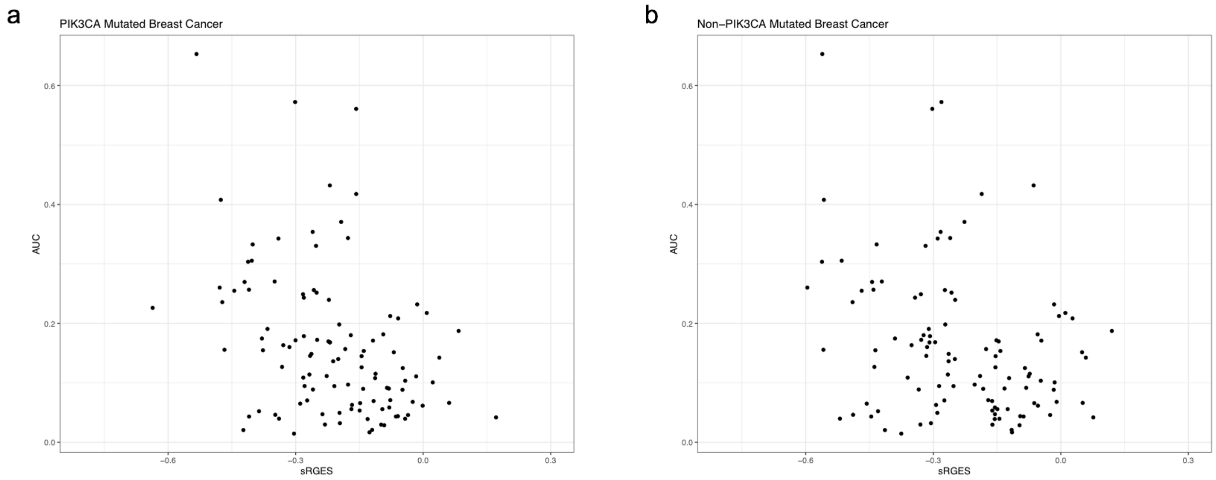


Figure S4: Correlation between sRGES and AUC in (a) PIK3CA mutated breast cancer, and (b) PIK3CA wild type breast cancer.
